# Wg/Wnt-signaling induced nuclear translocation of β-catenin is attenuated by a β-catenin peptide through its interaction with IFT-A in development and cancer cells

**DOI:** 10.1101/2023.06.14.544986

**Authors:** Linh T. Vuong, Marek Mlodzik

## Abstract

Wnt/Wingless (Wg) signaling is critical for many developmental patterning processes and linked to diseases, including cancer. Canonical Wnt-signaling is mediated by β-catenin, Armadillo/Arm in *Drosophila* transducing signal activation to a nuclear response. The IFT-A/Kinesin-2 complex is required to promote the nuclear translocation of β-catenin/Arm. Here, we define a small conserved N-terminal Arm/β-catenin (Arm^34–87^) peptide, which binds IFT140, as a dominant interference tool to attenuate Wg/Wnt-signaling *in vivo*. Expression of Arm^34–87^ is sufficient to antagonize endogenous Wnt/Wg-signaling activation resulting in marked reduction of Wg-signaling target gene expression. This effect is modulated by endogenous levels of Arm and IFT140, with the Arm^34–87^ effect being enhanced or suppressed, respectively. Arm^34–87^ thus inhibits Wg/Wnt-signaling by interfering with the nuclear translocation of endogenous Arm/β-catenin. Importantly, this mechanism is conserved in mammals with the equivalent β-catenin^34–87^ peptide blocking nuclear translocation and pathway activation, including in cancer cells. Our work indicates that Wnt-signaling can be regulated by a defined N-terminal peptide of Arm/β-catenin, and thus this might serve as an entry point for potential therapeutic applications to attenuate Wnt/β-catenin signaling.

## INTRODUCTION

Wnt/Wingless (Wg) signaling pathways play important roles in intercellular signaling in all metazoan organisms. They regulate many biological processes during embryonic development, including cell growth, cell migration, cell fate determination, cell polarity, stem cell homeostasis, and organogenesis in general^1–6^. Wnt-signaling pathways are evolutionarily conserved, and secreted Wnt/Wg proteins activate a variety of signal transduction events across all metazoans^4, 6, 7^. Wnt signaling is also one of the key pathways closely related to several diseases, including the initiation and progression of various types of cancers^8–13^.

The most studied Wnt-pathway, generally called “canonical Wnt-signaling”, is keyed by the nuclear translocation of β-catenin upon pathway activation. β-catenin, a.k.a. Armadillo (Arm) in *Drosophila*, is the “business-end” pathway component, with its cytoplasmic stabilization and nuclear translocation setting up its role as transcriptional co-activator, essential for Wnt-target gene expression^4–6, 14^. Wnt/Wg proteins bind to the Frizzled (Fz) and LRP5/6 (Arrow in *Drosophila*) co-receptors, and their binding results in the disassembly of the “destruction complex”, composed of Axin, APC (Adenomatous Polyposis Coli), and the kinases GSK3 and CK1. In the absence of Wnt ligands, the destruction complex recruits and phosphorylates the cytoplasmic pool of Arm/β-catenin and thus targets it for degradation^4–6,14–17^. Break-up of the destruction complex is mediated by Dishevelled (Dsh, Dvl in mammals), causing re-location of Axin to the plasma membrane, where it associates with Dsh/Dvl and the transmembrane Fz and LRP5/6 co-receptors. This leads to the formation of large complexes of Axin-Dsh-Fz-LRP5/6 aggregates, which are referred to as signalosomes^4, 6, 14, 18^. The removal of Axin from the destruction complex leads to stabilization of cytoplasmic Arm/β-catenin, which can then translocate to the nucleus to act as a co-activator of the TCF/LEF transcription factors^4, 6, 14, 15, 19, 20^.

In *Drosophila*, Arm/β-catenin plays roles at many different stages of development. Arm/β-catenin contains a large region formed by several repeats called the “Arm repeat domain” that is flanked by distinct N and C-terminal regions^21–23^. The Arm repeats of Arm/β-catenin form a concave groove region called an ARM domain that binds competitively to cadherins, APC, TCF/LEF1 and Axin^23–25^. While the Arm repeats domain plays a role in both canonical Wnt/Wg-signaling and the formation of adherens junctions (AJs)^23–25^, the C-terminal region has been shown to function exclusively for Wnt/Wg-signaling. It serves as an interaction surface for multiple complexes promoting Arm/β-catenin mediated transcription, via factors like CREB protein (CBP)/p300 and SET domain-containing protein 1 (SET-1)^21^, for example. Unlike the C-terminal region and the Arm repeat domain, the N-terminal region of Arm/β-catenin is less well understood, except for being critically required for its regulation by the destruction complex, coordinated by the scaffold protein Axin, and phosphorylation by CK1 and GSK3 kinases^26–28^. CK1 family members phosphorylate Arm/β-catenin at serine 45, required as priming phosphorylation for subsequent phosphorylation events by GSK3 at residues 41, 37, and 33 ^29^. In *Drosophila*, removal of the entire N-terminal Arm region or deletion of 53 amino acids (aa 34-87) around the phosphorylation sites (the latter called ArmS10) leads to highly stable cytoplasmic Arm/β-catenin protein and thus constitutive activation of Wnt/β-catenin signaling, independent of ligand activation^28, 30^.

Recently, we have shown that IFT-A forms a complex with Arm/β-catenin through the N-terminal region of Arm/β-catenin and that this interaction is important for its nuclear translocation^28, 31^. IFT-A (Intraflagellar Transport A Complex, known for its role in ciliogenesis) associates with kinesin 2, the *Drosophila* homolog of Kif3a and promotes nuclear translocation of Arm/β-catenin upon Wnt/Wg pathway activation^28, 31^. Loss of function of either IFT-A complex components, and IFT140 in particular, or Kinesin 2 proteins in *Drosophila* wing tissues displayed indistinguishable effects on Wnt/Wg signaling target, reflected by loss/reduction of their gene expression and associated wing development defects. Kinesin 2 directly interacts with IFT140 through Kap3, and acts as the motor to transport IFT-A along cytoplasmic microtubules. Both single and double mutant clones for *kinesin 2* and *ift140* fail to activate Wg/Wnt signaling targets in *Drosophila* ^28^. Moreover, double mutant clones for *IFT140* with *axin* displayed high levels of stabilized cytoplasmic Arm/β-catenin, in both wing imaginal disc cells and salivary gland cells, but target gene activation and its nuclear translocation were markedly reduced or even lost^28^. In this context, IFT140 directly binds to Arm/β-catenin through the N-terminal Arm^34–87^ region. It is thus an intriguing question to determine whether this N-terminal region of Arm/β-catenin, Arm^34–87^, plays a critical role in its nuclear translocation and how it might affect canonical Wnt-signaling.

In this study, we analyzed the mechanism of how this N-terminal peptide of Arm/β-catenin (Arm^34–, 87^) and IFT140, a component of IFT-A complex, function in canonical Wnt-signaling. We demonstrate that Arm^34–87^ specifically interacts with IFT140, both physically and genetically. Expression of this peptide is sufficient to antagonize endogenous Wnt/Wg-signaling activation. We demonstrate that this antagonism is mediated by competitive binding to IFT140, thus inhibiting the nuclear translocation of endogenous Arm/β-catenin. Importantly, this mechanism is conserved in mammalian cells. Our study defines an important role of the N-terminal region of Arm/β-catenin, and the specific peptide (residues 34-87) in particular, in Wnt/Wg-signal transduction, besides its known role in the destruction complex mediated process. It not only informs about a new mechanism but also provides insight into potentially therapeutic approaches and tools to attenuate canonical Wnt-signaling through (dominant) inhibition of its nuclear translocation and function.

## RESULTS AND DISCUSSION

### An Arm/β-catenin peptide, Arm^34–87^, is sufficient to bind to IFT140

We have shown previously that IFT-A and Arm/β-catenin are associated in a protein complex together with Kinesin 2, in which IFT140 directly interacted with Arm/β-catenin^28^. To confirm and further refine the interaction between IFT140 and Arm/β-catenin, we first established that the binding of a small protein fragment/peptide within the N-terminal domain of Arm, residues 34-87, is specific and sufficient to bind to IFT140 (Fig. 1A). In contrast, other components of the Kinesin 2 motor protein complex, Klp64D (*Drosophila* Kif3a) or Kap3 (Kinesin associated protein 3) did not bind to the Arm^34–87^ fragment, serving as control (Fig. 1A). To confirm these results *in vivo*, we performed co-immunoprecipitation (co-IP) assays with *Drosophila* wing disc extracts expressing Arm^34–87^-GFP and IFT140 or IFT144 (under *C96-Gal* control, which is expressed along the dorso-ventral wing disc region). Extracts from *C96>IFT140-myc; Arm*^34–87^*-GFP* and *C96>IFT144-myc; Arm*^34–87^*-GFP* wing discs were immunoprecipitated using anti-GFP and probed with anti-myc (Fig.1B), revealing that IFT140 co-IPed with Arm^34–87^, but IFT144 did not (Fig. 1B). Taken together, these data indicate that IFT140 directly binds to Arm/β-catenin within the small N-terminal region encompassing amino acids 34-87, hence called Arm^34–87^, and that this Arm peptide is sufficient for interaction with IFT-A via IFT140 both *in vitro* and *in vivo*.

**Figure 1:**
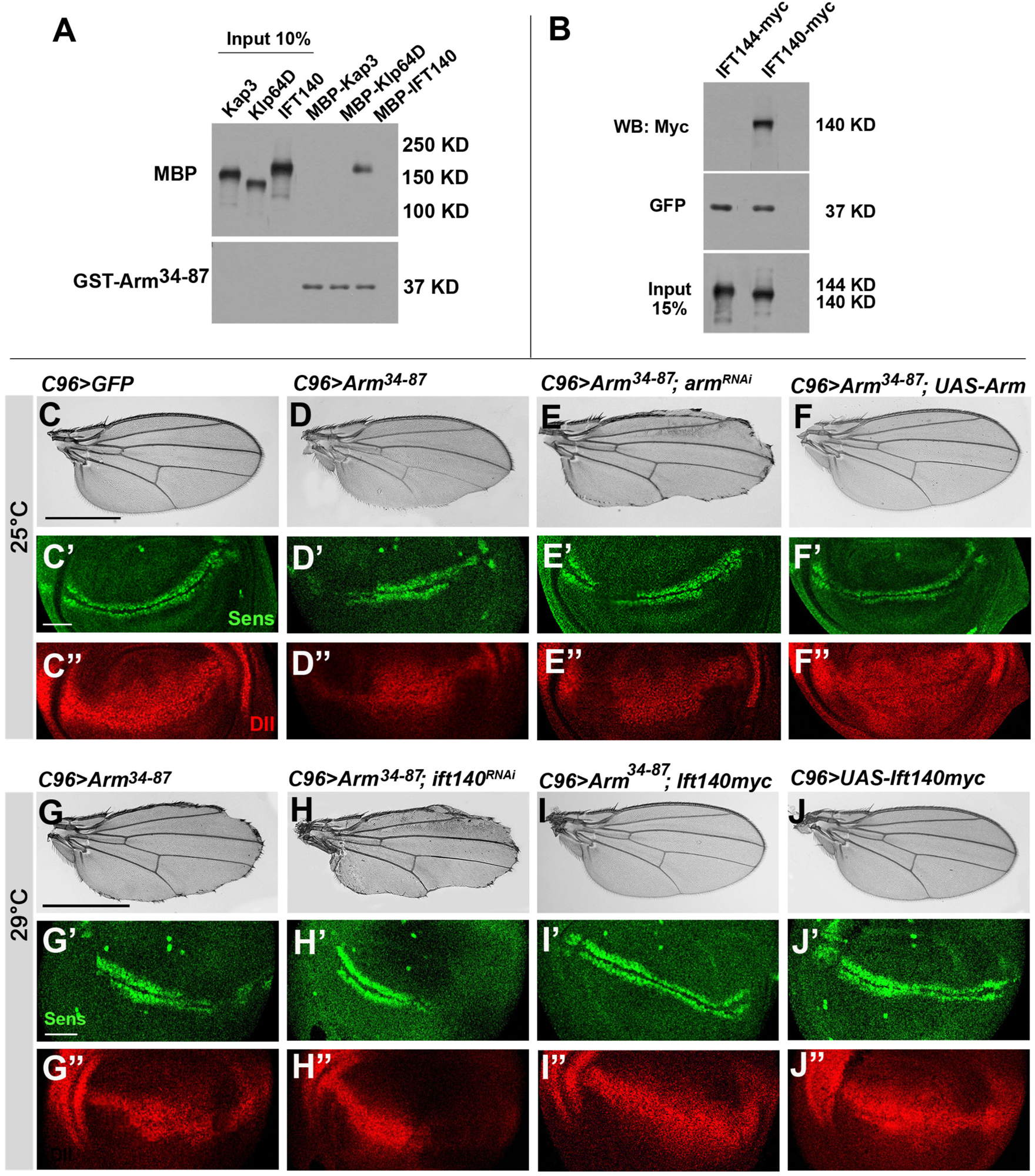
Arm^34–87^ peptide interferes with Arm/β-catenin signaling in *Drosophila* wing development. (**A-B**) Arm^34–87^ and IFT140 are physically associated. (**A**) Direct binding of Arm^34–87^ and IFT140. Full length Kap3, Klp64D (lane 1, 2: input 10%) were not pulled down by GST-Arm^34–87^ (lane 4, 5), but IFT140 (lane 3: input 10%) was pulled down by GST-Arm^34–87^ (lane 6). (**B**) *In vivo* co-immunoprecipitation (IP) assay of IFT144 and IFT140 with Arm^34–87^ from *tub>IFT144myc; Arm^34–87^GFP* (negative control, lane 1) and *tub>IFT140myc; Arm^34–87^GFP* (lane 2) wing imaginal discs. Protein extracts from wing discs were IPed with anti-GFP. IPs and input (15% of wing disc lysates used in IP step) were analyzed by blotting with myc antibody to IFT144 or IFT140. (**C-J”**) All wings and wing discs shown use the wing margin driver *C96-Gal4*. Adult wings: anterior is up and distal to right; wing discs: ventral is up and anterior to the left. (**C-F”**) Genetic interaction between *C96>Arm*^34–87^ and endogenous Arm. All genotypes were reared at 25C. (**C-C”**) *UAS-GFP* (*C96>GFP*) control wing, and control wing imaginal disc with wild-type Sens (green) (**C’**) and Dll (red) (**C”**) expression near D/V boundary. (**D-D”**) *C96>Arm*^34–87^ wing and wing disc. Note partial loss of margin (notching) phenotype (**D**) and partial loss of Sens (green, **D’**) and reduction of Dll (red, **D”**), consistent with adult wing defects. (**E-E”**) *C96>Arm*^34–87^*; >arm^RNAi^* (note that at 25C *>arm^RNAi^*alone shows no defects). Notching caused by Arm^34–87^ expression is enhanced, as well as Sens (**E’**) and Dll (**E”**) expression was further reduced (cf. to D-D”). (**F-F”**) *C96>Arm*^34–87^ *>Arm^wt^*: Note that notching caused by Arm^34–87^ is suppressed by co-expression of wild-type Arm, and Sens (green, **F’**) and Dll expression (red, **F”**) were restored. (**G-J”**) Genetic interaction between *IFT140* and *of Arm*^34–87^. All genotypes were reared at 29C. (**G-G”**) *C96>Arm*^34–87^: note increased notching with increased expression level at 29°C (cf. to panel D) and reduced Sens (**G’**) and Dll expression (**G”**). (**H-H”**) *C96>Arm*^34–87^*; >Ift140^RNAi^*: wing margin defects and Sens (**H’**) and Dll (**H”**) expression reduction were enhanced, compared to *C96>Arm*^34–87^ (cf. to G-G”). (**I-I”**) *C96>Arm*^34–87^*; >IFT140myc*: Co-expression of *IFT140myc* in the *C96>Arm*^34–, 87^ background suppressed wing notching phenotype and restored Sens (**I’**) and Dll (**I”**) expression. (**J-J”**) *C96>IFT140myc* control: note no effect of IFT140 by itself with wild-type margin and normal Sens and Dll expression in imaginal discs. The scale bar represents 100μm in adult wings (C and G) and 50μm in wing imaginal discs (C’-J”).

### Arm^34–87^ affects Arm/β-catenin function in Wg-signaling

Next, we wished to determine whether expression of the Arm^34–87^ peptide could impact Wg-signaling during development *in vivo*. To this end, we generated UAS-Arm^34–87^ and UAS-Arm^34–87^-GFP transgenes, both inserted at the same *attB*-chromosome site (59D3 on 2R, using the <C31 integrase technique to induce comparable expression levels)^32^. Ubiquitous expression of both, UAS-Arm^34–87^ and UAS-Arm^34–87^-GFP, caused larval lethality, suggesting that the N-terminal Arm peptide can dominantly interfere with normal development. To functionally define how Arm^34–87^ affects development we used *Drosophila* wings and wing margin patterning as model system. Strikingly, expression of UAS-Arm^34–87^ along the dorsal-ventral boundary region of wing discs, the future wing margin (under *C96-Gal4* driver control), resulted in partial loss of the margin (notching) (Fig. 1D, see Fig. 1C for control). Quantification of this notching phenotype, by measuring the length of margin loss, revealed that Arm^34–87^ expression resulted in 50% loss in margin fate. To ascertain whether Arm^34–87^ acted as a dominant-negative factor, interfering with endogenous Arm/β-catenin, we asked whether the phenotype of Arm^34–87^ expression can be modulated by reducing or increasing the levels of Arm/β-catenin. Indeed, the phenotypic effects of Arm^34–87^ expression were markedly enhanced by Arm knockdown via RNAi (Fig. 1E) or suppressed by co-expression of wild type full-length Arm/β-catenin (Fig. 1F).

*Distal-less* (*Dll*) and *senseless* (*sens*) are specific downstream transcription targets of Wnt/Wg-signaling during *Drosophila* wing development, with Sens being a high-threshold target, defining the future margin cells, and Dll a general wing specific target^33–36^. Consistent with the notion that Arm^34–87^ could interfere with canonical Wnt/Wg-signaling, *C96>Arm*^34–87^ expression leads to either loss of expression in a subset of cells along the wing margin (Sens) or markedly reduced expression in distal wing tissue (Dll) (Fig. 1D’-D”), as compared to control wing discs (Fig. 1C’-C”). Furthermore, defects in Sens and Dll expression induced by Arm^34–87^ expression were markedly enhanced or suppressed by Arm RNAi knockdown or full-length Arm/β-catenin co-expression (Fig. 1E’-E” and Fig. 1F’-F”’). To establish that the wing margin loss, notching phenotype induced by Arm^34–87^ expression was not caused by unspecific cell death, we asked whether the Arm^34–87^ phenotype can be suppressed by co-expressing the cell death inhibitor p35 (ref^37^). While wing notching defects caused by expression of the proapoptotic gene *hid* (*C96>UAS-hid*) was rescued by p35 co-expression (Suppl. Fig1), p35 co-expression had no noticeable effect on the Arm^34–87^ induced defects. Accordingly, Arm^34–87^ expression did not induce cleaved caspase 3 staining (Casp3) as a marker for cell death (Suppl. Fig1, note also that p35 efficiently suppressed cleaved Casp3 staining induced by Hid). Taken together, these data suggest that the dominant-negative *in vivo* effects of Arm^34–87^ during development and wing margin development were caused by inhibition of canonical Wnt/Wg-signaling at the level of Arm/β-catenin and Wg-signaling target gene expression.

### The Arm^34–87^ peptide interferes with IFT140 function

To determine whether the effects of Arm^34–87^ on canonical Wg-signaling were caused by its interaction with IFT140, and thus interference with IFT140/IFT-A function in canonical Wnt/Wg-signaling, we tested whether the phenotype of Arm^34–87^ expression was sensitive to IFT140 levels. Knock-down of *IFT140* (via RNAi or in mutant clones) displayed very similar notching to Arm^34–87^ expression^28, 31^. Strikingly, reducing levels of IFT140 enhanced the wing notching phenotype of Arm^34–87^, causing a significant increase of wing margin loss (Fig. 1H). In contrast, co-expression of wild-type IFT140 with Arm^34–87^, thus increasing IFT140 levels, suppressed the wing margin defects associated with Arm^34–87^ (Fig. 1I, cf. to Fig. 1G; note that *C96*-driven expression of IFT140 alone [in a wild-type background] had no effect on wing development, Fig. 1J). Consistent with the wing margin defects, expression of Arm^34–87^ analyzed in wing imaginal discs, revealed reduction or loss of Sens expression along the DV boundary, and reduction in Dll levels within the wing pouch area (Fig. 1G’-G”, as compared to the *wt* control, Fig. 1C’-C”). The Arm^34–87^ induced defects in Sens and Dll expression were considerably enhanced or rescued by *IFT140* RNAi co-expression (Fig. 1H’-H”) or IFT140 protein co-expression (Fig. 1I’-I”), respectively (note again that *C96>IFT140* alone caused no detectable defects in either Sens and Dll expression in wing discs; Fig. 1J’-J”). Taken the above results together, the data suggest that Arm^34–87^ is acting as a dominant-negative factor in canonical Wg-signaling by specifically interfering with the interaction between Arm/β-catenin and IFT140.

IFT140 and full-length Arm/β-catenin have been shown to co-localize in cytoplasmic punctae in *Drosophila* wing imaginal disc and salivary gland cells upon Wnt/Wg-signaling activation^28^. As Arm^34–87^ expression causes a dominant-negative effect on Wg-signaling and physically interacts with IFT140, we examined whether the Arm^34–87^ peptide co-localizes with IFT140 on its own and affects the overlapping localization of IFT140 and endogenous Arm/β-catenin. There are no specific antibodies against the Arm^34–87^ peptide, and so we used differentially tagged Arm^34–87^-GFP and full-length Arm-HA to distinguish between peptide and endogenous protein, together with IFT140-myc (staining with antibodies to the respective tags). To allow subcellular localization analyses of Arm with sufficient resolution, we used the *in vivo* Wg-signaling assay in the large cells of salivary glands^28^. In the absence of Wg expression, full-length Arm-HA was detected only at AJs (Fig. 2A, magenta), while Arm^34–87^-GFP (Fig 2A, green) displayed intracellular punctae. Interestingly, these punctae co-stained for IFT140-myc (Fig. 2A, red; also Suppl. Fig 2A), indicating (i) that the Arm^34–87^ peptide is not tethered to AJs, and (ii) that it can interact with IFT140 in the cytoplasm in the absence of Wg-signaling. Upon Wg expression, full length Arm was detected both at AJs and in cytoplasmic punctae (Fig. 2B; also^28^). Strikingly, some Arm-HA, Arm^34–, 87^-GFP and IFT140 triple positive punctae were detected (Fig. 2B, and Suppl. Fig 2A-C). These data indicate that the Arm^34–87^ peptide forms complexes with IFT-A independent of Wg/Wnt-signaling activation and thus its dominant negative behavior can be linked to this effect, in addition to competitive binding with endogenous Arm upon Wnt-signaling activation. The observed co-localization of Arm^34–87^-GFP and endogenous Arm upon Wg-signaling induction (Suppl. Fig 2B, C) was dependent on the presence of IFT140 (it was lost in *IFT140^cx2^* mutant allele; Suppl. Fig 2D), and thus dependent on IFT-A. This observation might suggest the formation of large complexes of several IFT-As.

**Figure 2.**
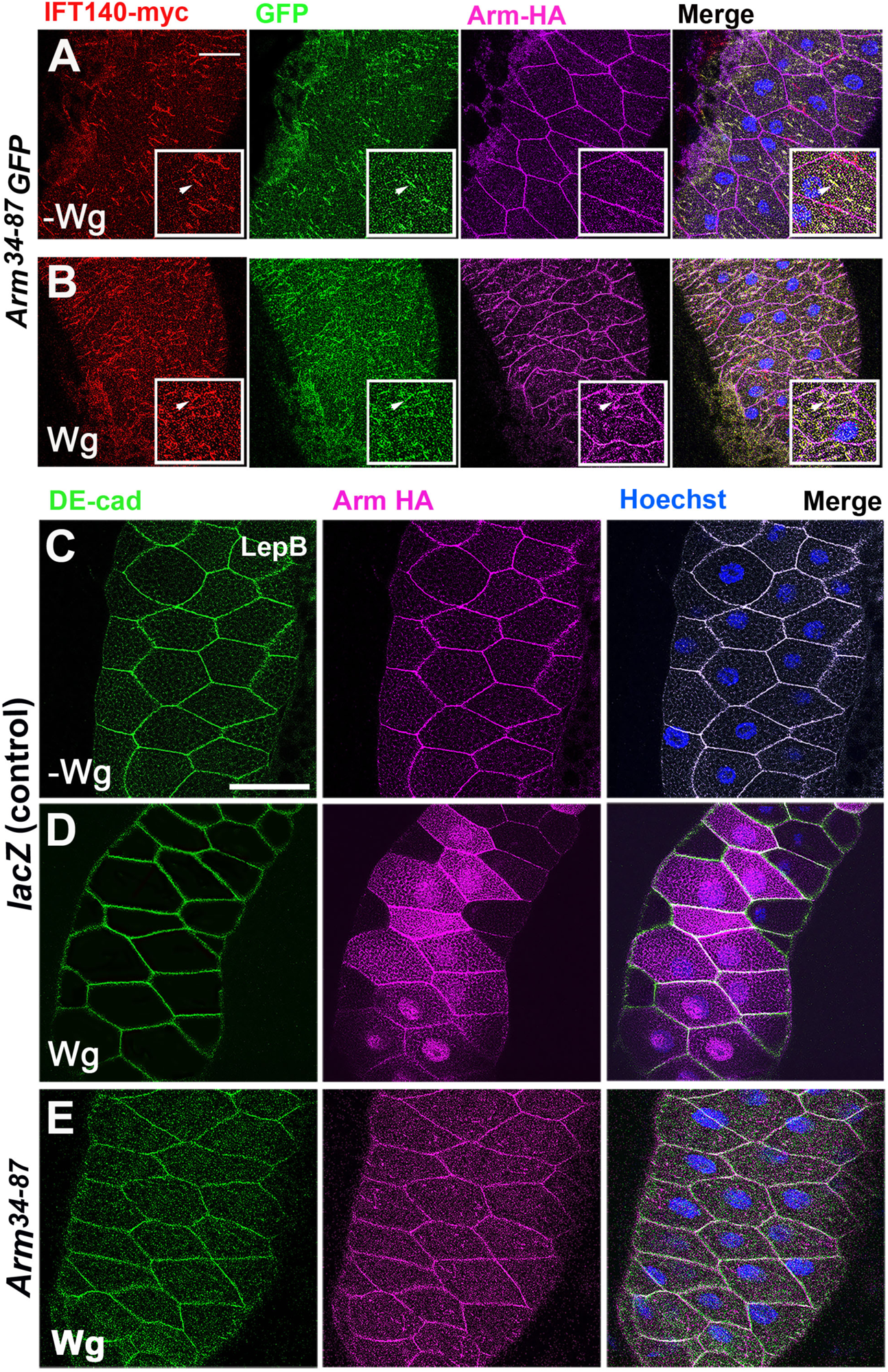
IFT140 and Arm^34–87^ co-localize in salivary gland cells and Arm^34–87^ blocks nuclear translocation of endogenous Arm/β-catenin. (**A-B**). Salivary glands were stained for IFT140-myc (red), Arm^34–87^-GFP (green, GFP), and Arm-HA (magenta). See inset for higher magnification. (**A**) Without Wg expression, Arm/β-catenin localizes to adherens junctions (AJs). Note that Arm^34–87^-GFP does not localize to AJs but displays punctate cytoplasmic staining overlapping with IFT140-myc. See inset for higher magnification and arrowheads marking co-stained puncta. (**B**) Upon Wg expression Arm/β-catenin is also detected in cytoplasm as it is stabilized and triple positive puncta staining for Arm^34–87^-GFP, IFT140-myc, and Arm-HA are detected in the cytoplasm. Inset shows higher magnification and arrowheads marking co-stained puncta. Scale bar is equal 50μm (**A-B**). (**C-E**) Wg-signaling induced Arm nuclear translocation assay. Salivary glands were exposed to Wg expression (via *C805-Gal4* driver, note mosaic expression of the driver, common to all SG gal4-drivers) and treated with Leptomycin B (LepB, which inhibits nuclear export) to enhance retention of Arm/β-catenin in the nucleus. Salivary glands were stained with DE-cad (green, membrane marker), Arm-HA (magenta), and Hoechst (nuclei, blue). Scale bar represents 100μm. (C) *UAS-lacZ* (without expression of Wg, control), Arm/β-catenin mainly localizes to AJs at the membrane. (**D**) Wg-expression (*>Wg,>LacZ*; positive control): note Arm/β-catenin localization in both cytoplasm and nucleus (uneven increase in Arm/β-catenin levels is caused by mosaic expression of Gal4-driver). (**E**) Co-expression of Arm^34–87^ peptide with Wg (*>Wg, >Arm*^34–87^) largely eliminates nuclear translocation of endogenous Arm/β-catenin with increased punctate localization in the cytoplasm.

Overall taken together, these results suggest (i) that Arm^34–87^GFP is purely cytoplasmic and does not associate with Ajs, (ii) that the majority of Arm^34–87^GFP co-localized with IFT140 in the presence or absence of Wg signaling activation, and (iii) that Arm^34–87^GFP and Arm-HA can display overlapping cytoplasmic localization, which is dependent on IFT140, and this in turn suggests that these are large complexes of unknown stoichiometry.

### Arm^34–87^ inhibits nuclear localization of endogenous Arm/β-catenin

We have previously demonstrated that the Kinesin 2/IFT-A complex is required for nuclear localization of Arm/β-catenin ^28^. Since the Arm^34–87^ peptide is necessary and sufficient for binding of Arm/β-catenin to the kinesin-2/IFT-A complex (via its interaction with IFT140), we next examined whether Arm^34–87^ could affect nuclear translocation of endogenous Arm upon Wg-signaling activation, using the established *in vivo* assay system in the salivary glands. Wg-signaling was activated via the salivary gland (SG) specific *C805-Gal4* driver expression in SG cells (note that this and all other SG-drivers display mosaic expression), inducing nuclear translocation of Arm/β-catenin (detected with anti-Arm staining; to facilitate detection of nuclear Arm, SGs were treated with Leptomycin B before fixation)^38^. *Drosophila* E-cadherin (DE-cad) and Hoechst were used as cell membrane and nuclear markers, respectively. Expression of Wg activated the pathway and revealed increased levels of Arm, detected both in nuclei and cytoplasm, demonstrating the Wg-induced Arm/β-catenin stabilization and nuclear translocation (Fig. 2D, note that the cytoplasmic Arm stabilization and nuclear level was variable from cell to cell due to the mosaic expression of the *Gal4*-driver). In contrast, Arm/β-catenin was only detected at AJs, when Wg was not expressed and hence Wg-signaling not activated (Fig. 2C). Moreover, co-expression of Arm^34–87^ with Wg in SGs revealed a striking loss of nuclear Arm/β-catenin staining (Fig. 2E), indicating that Arm^34–87^ was interfering with nuclear translocation of endogenous Arm/β-catenin.

Taken together with the genetic and biochemical results involving IFT140, these data strongly suggest that the Arm^34–87^ peptide can interfere with the Kinesin2/IFT-A complex mediated nuclear translocation of Arm/β-catenin by competing for binding to that complex and thus attenuating endogenous Wnt-signaling by blocking endogenous Arm/β-catenin access to the nucleus.

### Function of the Arm/β-catenin^34–87^ peptide is evolutionarily conserved

The sequence across the Arm^34–87^ region is fully conserved in β-catenin of higher animals^28^. To determine whether its function, being required for and capable of inhibition of nuclear localization of β-catenin, is conserved in mammals, we tested the respective mouse β-catenin peptide in mouse embryonic fibroblasts (MEFs). Wild-type (*wt*) MEFs revealed detectable levels of nuclear β-catenin upon Wnt3a stimulation (Fig. 3A; also^28^) and the same behavior was observed with *wt*-MEFs transfected with full length β-catenin-GFP (Fig. 3E). In contrast, under the same Wnt3a treatment conditions, the β-catenin^34–87^-GFP peptide transfected *wt*-MEFs displayed a loss of endogenous nuclear β-catenin (Fig. 3C, G; Suppl. Fig 3; note that β-catenin^34–87^-GFP alone also stayed in the cytoplasm, Fig. 3B, like its *Drosophila* counterpart, see above), indicating a conserved behavior of the β-catenin^34–87^ peptide in blocking nuclear translocation of endogenous full-length β-catenin. Importantly, and consistently with the experiments in *Drosophila*, this function was dependent on Wnt-signaling induction, as without Wnt3a treatment neither displayed pathway activation and nuclear β-catenin (Fig. 3B, D, F, and H).

**Figure 3.**
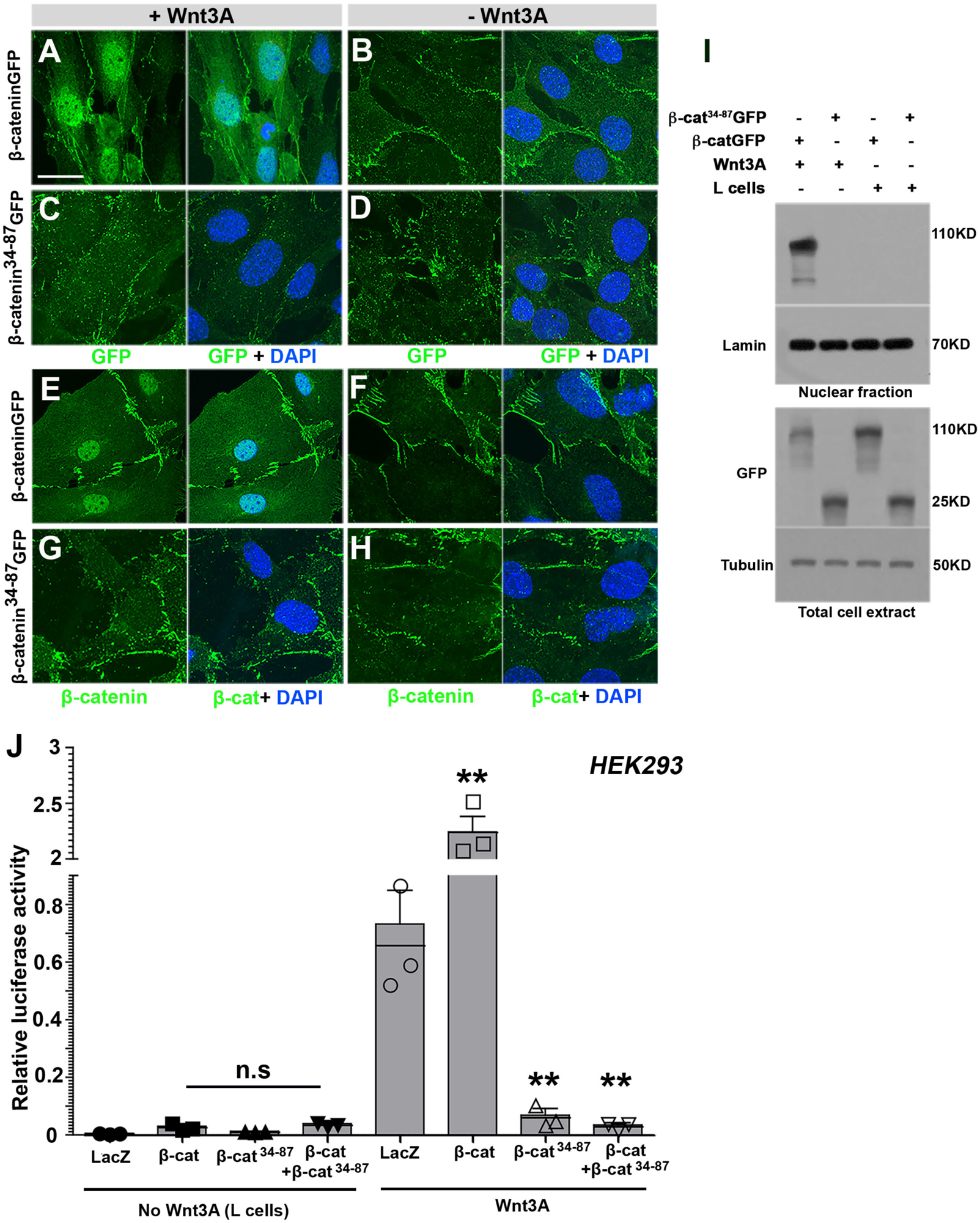
β-catenin^34–87^ peptide inhibits nuclear accumulation of β-catenin in mammalian cells MEFs and inhibits Wnt signaling target expression. (**A-H**) Confocal images of immunofluorescence (IF) of β-catenin or the β-catenin^34–87^ peptide (green, single channel of left) and nuclei (blue in overlay) in MEFs treated with Wnt3a-conditioned (**A, C, E, G**) or unconditioned control media (**B, D, F, H**). Upon 24h of Wnt3A stimulation, β-catenin was readily detectable in nuclei of wild-type MEFs either by GFP antibody (**A**) or β-catenin antibody (**E**); in contrast, it was undetectable in nuclei of MEFs with β-catenin^34–87^ transfection, either by GFP (**C**) or β-catenin antibody (**G**). No nuclear β-catenin was detected in any MEFs cultured with unconditioned media (B, D, F, H: controls). IF signal of membrane associated β-catenin was undistinguishable in all MEFs, cultured with Wnt3A-conditioned or unconditioned media (A-H). Scale bar represents 50μm. **(I)** Western blots of nuclear and total cell fractions with treatment of Wnt3A or without (negative control media: L-cells supernatant). γ-tubulin and Lamin B were used as loading controls for cytoplasm and nuclei, respectively. Note nuclear accumulation of full-length β-catenin upon Wnt-treatment and the lack thereof in the presence of β-catenin^34–87^-GFP. **(J)** β-catenin^34–87^ peptide inhibits Wnt-signaling target gene activation in human HEK293 cells. Wnt-signaling activity in HEK293 cells was assayed by comparing Wnt3a stimulated cells and control cells transfected with either *LacZ* (control), β-catenin, β-catenin^34–87^ peptide, or β-catenin and the β-catenin^34–87^ peptide, respectively (as marked on X-axis in graph). Relative luciferase activity (Y-axis) shows the ratio of firefly TOP-Flash reporter and renilla luciferase control. Wnt3A conditioned media were added 48 hrs prior to cell lysis to induce signaling. Overexpression of β-catenin increases luciferase activity, whereas transfection of the β-catenin^34–87^ peptide causes a strong reduction of Wnt-signaling activity (\*\**p* <0.001 compared to LacZ control, three independent assays).

To confirm these observations, we also analyzed nuclear β-catenin levels by Western blotting in these backgrounds. While control *wt*-MEFs displayed a strong accumulation of nuclear β-catenin upon Wnt3a treatment (Fig. 3I), MEFs transfected with the β-catenin^34–87^-GFP peptide displayed hardly any detectable nuclear β-catenin under the same Wnt3a treatment (Fig. 3I). In summary, these data indicated that the N-terminal β-catenin^34–87^ peptide displayed the same behavior as *Drosophila* Arm^34–87^, with both inhibiting nuclear localization of Arm/β-catenin upon Wnt/Wg signaling activation.

### The β-catenin^34–87^ peptide inhibits Wnt signaling in human cancer cells

The dominant-negative effects of Arm/β-catenin^34–87^ in *Drosophila* wing margin development and the above effects on nuclear β-catenin translocation in MEFs, suggested that it can function as an inhibitor of Wnt-signaling in general, and thus also in human Wnt-signaling contexts. To address this, we carried out Wnt-pathway activation assays using the Top-Flash reporter (a luciferase reporter directly under TCF/μ-catenin DNA binding and transcriptional activation control) to measure target gene activation directly in human cells. We first assayed this in HEK293 cells (human embryonic kidney cells) transfected with LacZ as control in the absence or presence of Wnt in the medium, with Wnt3A-containing medium being added two days after transfection to induce signaling. Without Wnt3A addition, reporter activation (measured as the ratio of TCF-responsive luciferase reporter [TOP-Flash] and constitutive Renilla luciferase expression) was near zero. However, with addition of Wnt3A, the relative luciferase activity increased significantly (Fig. 3J), and the activity was further increased by overexpressing exogenous β-catenin (Fig. 3J). Conversely, β-catenin^34–87^ peptide expression inhibited Wnt signaling to almost the basal level (compare to no Wnt3a medium addition in Fig. 3J). When β-catenin and the β-catenin^34–87^ peptide were co-expressed, Wnt signaling activity was markedly suppressed below its endogenous levels, even in the presence of overexpressed full-length β-catenin (Fig. 3J). These results indicate that the β-catenin^34–87^ peptide strongly inhibits Wnt-signaling target gene activation in human cells.

To further investigate the inhibitory potential of the β-catenin^34–87^ peptide on Wnt/β-catenin signaling in disease contexts, we tested its effect in human cancer cells. To this end, we assessed its impact on Wnt signaling in lung cancer cell lines (A549, H2009 and H1299), breast cancer cell line HCC1395, and brain tumor line SF25. Western blot analysis was used to determine endogenous Wnt3A levels in these cancer cell lines (Fig 4F and Suppl. Fig 4A). Wnt3A was expressed at high levels in HCC1395, but not in the A549 cell line (Fig. 4F). We thus first asked whether in the A549 cell line addition of Wnt3a could induce endogenous nuclear β-catenin localization, which is indeed the case (Fig. 4A, B). Strikingly, co-transfection of the N-terminal β-catenin^34–87^ peptide blocked the nuclear localization of endogenous β-catenin itself (Fig. 4B). Consistently, it also markedly reduced Wnt3a-dependent gene expression as measured in the TOP-Flash assay (Fig. 4C) and proliferation in the A549 line (Fig. 4D, E). Very similar to the A549 cell line, the luciferase reporter system activated by β-catenin/TCF (ref^39^) was suppressed by transfection of the β-catenin^34–87^ peptide in H2009 and H1299 lung cancer cell lines and the SF295 brain tumor line (Suppl. Fig 4B-D; note that these cancer lines express Wnt3a endogenously in an autocrine manner, Suppl. Fig. 4). Next, we analyzed the breast cancer line HCC1395, which also expresses Wnt3A endogenously. Very similar to the other cancer lines, transfection of the β-catenin^34–87^ peptide markedly reduced proliferation of HCC1395 cells (Fig. 4H, I) and almost completely eliminated activation of the Wnt-signaling β-catenin/TCF-reporter (Fig. 4G).

**Figure 4.**
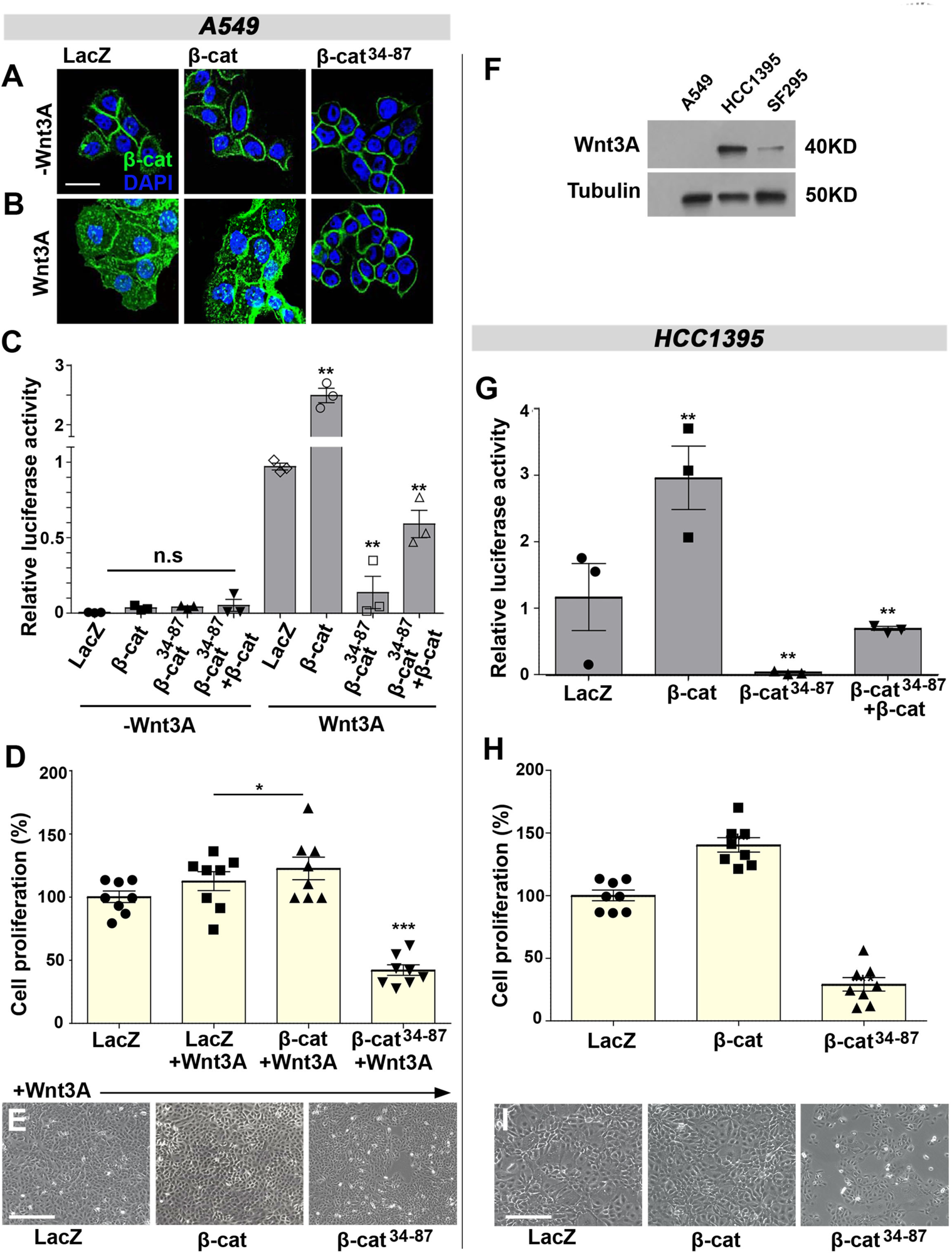
The β-catenin^34–87^ peptide inhibits Wnt signaling and cell proliferation in human cancer cell lines. (**A-E**) A549 lung cancer cells were transfected with LacZ, β-catenin-GFP or β-catenin^34–87^-GFP with or without Wnt3A conditioned media. (**A-B**) Cells were stained with anti-β-catenin (green) and DAPI (blue, nuclear marker). LacZ, β-catenin-GFP and β-catenin^34–87^-GFP transfected cells showed β-catenin localization at cell-cell junctions without Wnt3A treatment (**A**). Upon treatment with Wnt3A, punctate β-catenin staining is detected in both cytoplasm and nucleus (**B**, left and middle panels), except when co-transfected with β-catenin^34–87^-GFP peptide, which eliminated nuclear β-catenin translocation with Wnt3A treatment (**B**, right panel). Scale bar represents 200μm (**C**) Relative luciferase activity assay in A549 cells. To characterize the β-catenin^34–87^ effect on Wnt/β-catenin signaling activity, TOP-Flash firefly Luciferase reagent was added to cells first and then Renilla Luciferase reagent was added. Co-transfection of *wt* β-catenin increases the relative luciferase activity compared to LacZ control. In contrast, transfection of the β-catenin^34–87^ peptide markedly reduced reporter activity in the absence or presence of *wt* β-catenin (** *p* <0.001, three independent assays). (**D-E**) Cell proliferation assay in A549 cells. Cells were transfected with LacZ, β-catenin-GFP or β-catenin^34–87^-GFP together with Wnt3A (except control). Note significant drop in cell proliferation (quantif. in **D;** \**p* <0.05 and \*\*\**p* <0.0001, three independent assays), as well as detected by cell density in photographs (**E**, taken at 72hr after Wnt treatment). Scale bar represents 500μm. **(F)** Analyses of endogenous Wnt3A expression in several cancer cell lines (A549, HCC1395 and SF295). Note high levels of Wnt3a in HCC1395 cells. (**G-I**) β-catenin^34–87^-GFP blocks Wnt-signaling and reduces proliferation in HCC1395 cancer cells. **(G)** Relative luciferase assay as described in panel C. Note that co-transfection of *wt* β-catenin increases the relative luciferase activity (compared to LacZ control) and that transfection of the β-catenin^34–87^ peptide markedly reduces relative reporter activity in the absence or presence of *wt* β-catenin (** *p* <0.001, three independent assays). (**H-I**) Proliferation of HCC1395 cells is suppressed by β-catenin^34–87^ peptide. Cells were transfected with LacZ (control), β-catenin-GFP or β-catenin^34–87^-GFP. After 3 days, proliferation was assessed via the cell proliferation assay (see Methods). Note markedly reduced proliferation in β-catenin^34–87^-GFP transfected cells (quantif. in **H**; \*\*\**p* <0.0001, three independent assays) and also visible as cell density in growing cells (**I)**, with photographs taken at 72hrs after transfection. Scale bar represents 500μm.

Taken together with the mechanistic studies in the previous sections, these results strongly suggest that the β-catenin^34–87^ peptide may be effective in reducing or even blocking Wnt-signaling in a multitude of tumors.

## CONCLUSIONS

We have here defined a specific peptide in the N-terminal region of Arm/β-catenin, residues 34-87, hence referred to as Arm/β-catenin^34–87^, as sufficient for its physical interaction with IFT140, a component of the IFT-A complex. We have shown previously that the Kinesin 2/IFT-A complex is required for nuclear translocation of Arm/β-catenin (ref^28^). Expression of Arm^34–87^ in the DV boundary region giving rise to the wing margin of *Drosophila* wings causes Wg-signaling loss-of-function defects, manifest in wing margin loss (a.k.a. notching) and associated loss and reduction of the expression of the Wg-signaling targets Sens and Dll. In the *in vivo* salivary gland Wg-signaling assay Arm^34–87^ co-expression inhibits nuclear translocation of endogenous Arm/β-catenin and the same inhibitory behavior on Wnt-signaling is observed in mammalian contexts, with β-catenin^34–87^ transfection markedly blocking Wnt3a-induced nuclear translocation of endogenous β-catenin and associated reduction in Wnt-target gene activation in MEFs, HEK293 cells, and several human cancer cell lines. In addition, ubiquitous Arm^34–87^ expression during *Drosophila* embryogenesis is lethal during early development. Taken together with the fact that it is sufficient for IFT-A interaction and that it does not appear to affect junctional Arm/β-catenin, we conclude that it generally exerts all its effects by blocking Wnt-induced nuclear translocation of endogenous Arm/β-catenin and associated loss/reduction in Wnt-target gene activation.

Arm/β-catenin is a multifunctional protein associated with cell adhesion at AJs, linking these to the actin cytoskeleton, and canonical Wnt signaling, where it acts as the key nuclear effector. The entry of cytoplasmic Arm/β-catenin into the nucleus is a critical event in Wnt/β-catenin signaling. Since Arm/β-catenin has no nuclear localization sequence (NLS), it has long been speculated on how it gets translocated into the nucleus to promote transcription as a co-activator^40^. Recently, we have identified a requirement of the Kinesin 2/IFT-A complex for nuclear translocation of Arm/β-catenin ^28^, with IFT140 (within the IFT-A complex) and Arm/β-catenin directly interacting through a small region within the N-terminus of Arm/β-catenin (Arm^34–87^). Deletion of these residues results in a stable isoform, called ArmS10 (ref^30^), as it also contains all phosphorylation sites targeted by the destruction complex. ArmS10 is thus commonly used in *Drosophila* as a “constitutively active” Arm/β-catenin isoform^30^. In fact, ArmS10 can enter the nucleus independently of the Kinesin 2/IFT-A function^28^. While this was surprising at first, Kinesin 2/IFT140 independent nuclear translocation mechanisms have been recently proposed as well^41^ and ArmS10 does not require further protection from association with the destruction complex due to the deletion. The data presented here argue strongly that Kinesin 2/IFT140 dependent nuclear translocation is the primary mechanism for the process, as functional studies with the Arm/β-catenin^34–87^ peptide markedly reduced nuclear β-catenin in both *Drosophila* and mammalian cells.

The Arm^34–87^ peptide is necessary and sufficient to bind to IFT140 within the IFT-A complex. Genetic and functional *in vivo* assays suggest that the effect of Arm/β-catenin^34–87^ expression is mediated by a dominant-negative behavior on the IFT-A interaction of endogenous Arm/β-catenin protein. This model is supported by the suppression or enhancement of Arm^34–87^ wing notching phenotypes by co-expressing or silencing IFT140, respectively. Similarly, an increase of full-length Arm/β-catenin levels can suppress the dominant effect of the N-terminal peptide, largely rescuing the wing phenotype and Sens and Dll expression back to wild-type. Taken together, our biochemical data and co-localization studies suggest that its dominant effect is mediated by competitive binding between the Arm/β-catenin^34–87^ peptide and endogenous full-length Arm/β-catenin to IFT140. Our results also suggest that the Arm^34–87^ fragment is functionally critical for the interaction between IFT140 and Arm/β-catenin, and that this domain is essential for normal function of Arm/β-catenin in nuclear translocation during development and disease.

With β-catenin being the “business-end” of canonical Wnt-signaling, the key effector responsive for signal transduction to the nucleus, our data also suggest that nuclear translocation might be explorable as a therapeutic target to attenuate an overactivated Wnt-response. We show here that Wnt-addicted (autocrine) cancer cells can be inhibited by expression of the Arm/β-catenin^34–87^ peptide. This feature could also be explored at the level of diagnostics, as cancer cells “self-activating” via autocrine Wnt-signaling are detectable by the β-catenin^34–87^ peptide treatment assay. We thus not only define the function of a small region within the N-terminal domain of Arm/β-catenin, but our data should serve as an entry point for potential new diagnostics, and/or drug development and therapeutic applications to detect and inhibit overactive Wnt/β-catenin signaling in disease contexts, including cancer. This is of particular significance, as there are currently no approved drugs that inhibit canonical Wnt signaling at the level of β-catenin or at any level of the signaling pathway.

## MATERIAL AND METHODS

### Fly stocks

The Gal4/UAS system was used for expression of RNAi constructs (sometimes in combination with *UAS-Dcr2*) and other transgenes. Gal4-driver for wing margin during wing development was *C96-Gal4* expressed around the dorsal-ventral compartment boundary of wing imaginal discs, and the Gal4-driver for salivary glands was *C805-Gal4.* All crosses were set up at 25°C or at 29°C, as indicated.

### MARCM mutant clones

To generate *arm* mutant clones, males of *hs-FLP, FRT 19A tub-Gal80; nub-Gal4, UAS-GFP/UAS-GFP* were crossed with females *FRT19A arm*^2^*/FM7c* flies. Control clones were generated by crossing males of *hs-FLP, FRT19A, Tub-Gal80; nub-Gal4, UAS-GFP/UAS-GFP* with females of *FRT19A/FRT19A*. Mutant clones were induced at 60h after egg laying (AEL) by one hour heat shock at 37°C. Larvae were aged at room temperature (RT) until 3^rd^ instar (about 120h AEL) and wing imaginal discs were dissected for immunostaining.

### Immunostaining and histology

Imaginal discs were dissected at 3^rd^ instar larval stage in PBS and fixed in PBS, 4%PFA. Discs were washed 2 times in PBS 0.1% Triton-X100 (PBT), incubated in primary antibodies o/n at 4°C. After washing in PBT, incubation with secondary antibodies was at RT for 2hrs. Samples were mounted in Vectashield (Vector Laboratories). Wing disc images were acquired with a confocal microscope (20X-40X, oil immersion, Leica SP8 or Zeiss LSM880 system). Images were processed with ImageJ (National Institutes of Health) and assembled in Photoshop (Adobe).

Salivary glands were dissected at 3^rd^ instar larval stage in PBS and treated with 0.1% Leptomycin B (Sigma) in 10 min before fixation in PBS, 4% PFA. All subsequent steps were as described for wing imaginal discs.

Analyses of adult wings: wings were removed, incubated in PVT, and mounted on a slide in 80% glycerol in PBS, and imaged using Zeiss Axioplan microscope. All adult images were acquired using Zeiss Axiocam color-type 412-312 camera and the Zeiss axiocam Zen software.

### Transgene construction

To generate transgenic flies, Arm^34–87^ and Arm^34–87^ were amplified by PCR using DGRC LD23131 cDNA (for Arm) and cloned into *pUAS-attB* and *pUAS-attB-GFP* vectors (VK1, second chromosome 2R 59D3) using NotI and XbaI sites. The following primers were used to make Arm^34–, 87^ constructs:

AR^34–87^: 5’-GCGGCCGCCAAAATGGGAGATGGAGGGAGATCCACT - 3’ and 5’-GCTCTAGACATACCGGTGTCCAGGTCGAA - 3’

### GST pull down

For GST pull-downs, IPTG-inducible E. coli R2 cells (BL21) were transformed with plasmid constructs for fusion proteins MBP-Kap3, MBP-Klp64D, MBP-IFT140 and GST-Arm^34–87^. Bacterial lysates were prepared as describe in. An equal amount of blocked glutathione Sepharose 4B beads (Bioprogen) with GST, GST fusion protein or beads alone were incubated with lysates containing MBP-fusion proteins O/N at 4°C. After several washes with pull-down buffer (20 mM Tris pH 7.5, 150 mM NaCl, 0.5 mM EDTA, 10% glycerol, 0.1% Triton X 100, 1mM DTT, and protease inhibitor cocktail), sample buffer was added, beads were boiled, and protein were resolved by SDS-PAGE. For Western blotting, proteins were transferred onto nitrocellulose, blocked in 5% skim milk (Biorad) and incubated with primary goat anti-GST (Santa Cruz) or rabbit anti-MBP antibody (Santa Cruz). Protein bands were visualized using ECL kit (Millipore).

### Immunoprecipitation

Lysates from 30 wing imaginal discs of *C96>IFT140myc, Arm^34–87^GFP*, *C96> IFT144myc,* Arm^34–^*^87^GFP* were precleared by incubating with protein A-sepharose beads (Thermo Scientific) for 1hr at 4°C followed by centrifugation. A-sepharose beads were immune-precipitated with specific antibodies at 4°C for 1hr. Polyclonal anti-GFP antibody (Roche) was used. Immunoprecipitates were resuspended in SDS sample buffer, boiled for 5 min, separated by SDS-PAGE, and transferred to nitrocellulose for immunoblotting. Protein was detected by enhanced chemiluminescence (Millipore).

### MEF immunofluorescence staining and Wnt3a-induced β-catenin localization

For Wnt3A-induced β-catenin nuclear translocation: MEF were grown at 70% confluence in DMEM medium supplemented with 10% fetal bovine serum and treated with Wnt3A-conditioned medium, for 14-16h. Cells were then fixed in cold methanol for 15 min at -20°C and labeled with primary antibodies for GFP or β-catenin diluted 1:100 in 2% bovine serum albumin/PBS for 2 hrs, washed in PBS, incubated with fluorescent secondary antibodies (AlexaFlour 488, Life Technologies, Carlsbad) diluted 1:500 in PBS for 1 hr, and mounted with Vectashield mounting medium.

### Quantification of β-catenin nuclear translocation via microscopy analysis

Confocal imaging was performed using a Leica SP8 microscope. Z stacks of 6 (1.083µm) to 12 optical slices (4.362µm) at 8-bit were captured and analyzed via ImageJ-Fiji. β-catenin translocation was evaluated through optical density assays of mean gray values between the nucleus and cytoplasm of individual cells. To maintain uniformity in all acquisitions, measurements were obtained under an image size of 1024×1024 pixels, FITC (green) channel, maximum projection (Z-project on Image J), and a standardized region of interest at 4.705µm x 4.843µm avoiding cell membrane regions. Intensity ratios (nucleus/cytoplasm), standard deviations, and student’s two-tailed test were performed using Microsoft Excel.

### β-catenin nuclear fraction assay

Wnt3A stimulated or unstimulated MEFs were gently washed with PBS and cells were minced on ice by sharp scalpel and collected by centrifugation at 5000rpm/4°C. Samples were resuspended in 500µm buffer comprising 250mM sucrose, 50mM Tris-Cl pH7.4, 5mM MgCl2, 1M EDTA, 1% TritonX100 and protease inhibitor cocktail (Roche) and gently homogenized for 1 min on ice using homogenizer. Samples were kept for 30 min at 4°C (on ice) in microfuge tubes, and supernatants were cleared by 20 min centrifugation at 4°C and saved as the cytoplasmic fraction. The pellet, or nuclear fraction, was washed by the lysis buffer without protease inhibitor cocktail. Nuclear fraction was incubated for 30min at 4°C with nuclear extract buffer containing 20mM HEPES ph7.9, 15mM MgCl2, 0.5M NaCl, 1M EDTA, 20% glycerol, 1% TritonX100 protease inhibitor cocktail and sonicated for 10” after incubation. The resulting supernatant was collected by 30min centrifugation at 4°C as the nuclear fraction. Protein extracts were boiled for 5min at 95°C in SDS-sample buffer, separated by 10% SDS-page gel and transferred to nitrocellulose. Protein levels were analyzed by immunoblotting with the corresponding antibodies.

For β-catenin isoform experiments: β-catenin^34–87^GFP construct was generated from the original plasmid MSCV-β-catenin-IRES-GFP (Addgene Plasmid #14717). These constructs were transfected into MEFs by Lipofectamine LTX&Plus^TM^ (Invitrogene A12621). Cells were collected 12 hrs after transfection.

### Cell lines and culture conditions

HEK293 cells were grown in Dulbecco’s Modified Eagle’s Medium (DMEM) (Sigma-Aldrich, St. Louis, MO) supplemented with 10% fetal bovine serum (FBS) (Sigma-Aldrich), 50 units/ml of penicillin/streptomycin. A549, H2009, H1229, HCC1395 and SF295 human cancer cell lines were grown in RPMI-1640 (Thermo Fisher Scientific, Waltham, MA) medium supplemented with 10% FBS and 50 units/ml of penicillin/streptomycin.

### Luciferase reporter assay for β-catenin activity

TOP-FLASH luciferase reporter vectors (Upstate Biotechnology, Lake Placid, NY) were used to measure β-catenin signaling activity. A549, H2009, H1229, HCC1395 and SF295 cells were seeded into 6 well plates and transfected with 0.3 μg TOPflash (containing wild-type TCF binding sites) negative control LacZ (20 MOI) in serum free medium. After 12hr, the medium was replaced with 1% DMEM and the cells were incubated for another 24 hr. Cells were lysated with passive lysis buffer, and 20 μl of the cell extract was analyzed using Dual-Luciferase Reporter Assay system (Promega, Madison, WI). Each experiment was carried out in triplicate, and repeated at least three independent times.

### Cell proliferation assay

The cell proliferation assay was determined by 3-(4, 5-dimethylthiazol-2-yl)-2,5-diphenyl-tetrazolium bromide (MTT) assay (Sigma). All cancer cell lines were seeded in 24-well plates (2×10^4^ cells/well). After 24hr, cells were transfected with lacZ, β-catenin or the β-catenin^34–87^ peptide. The following day, cells were stimulated with or without recombinant Wnt3A (100ng/ml) for an 48hr. Absorbance at 540nm was read on a microplate reader. All assays were performed in triplicate.

## KEY RESOURCES TABLE

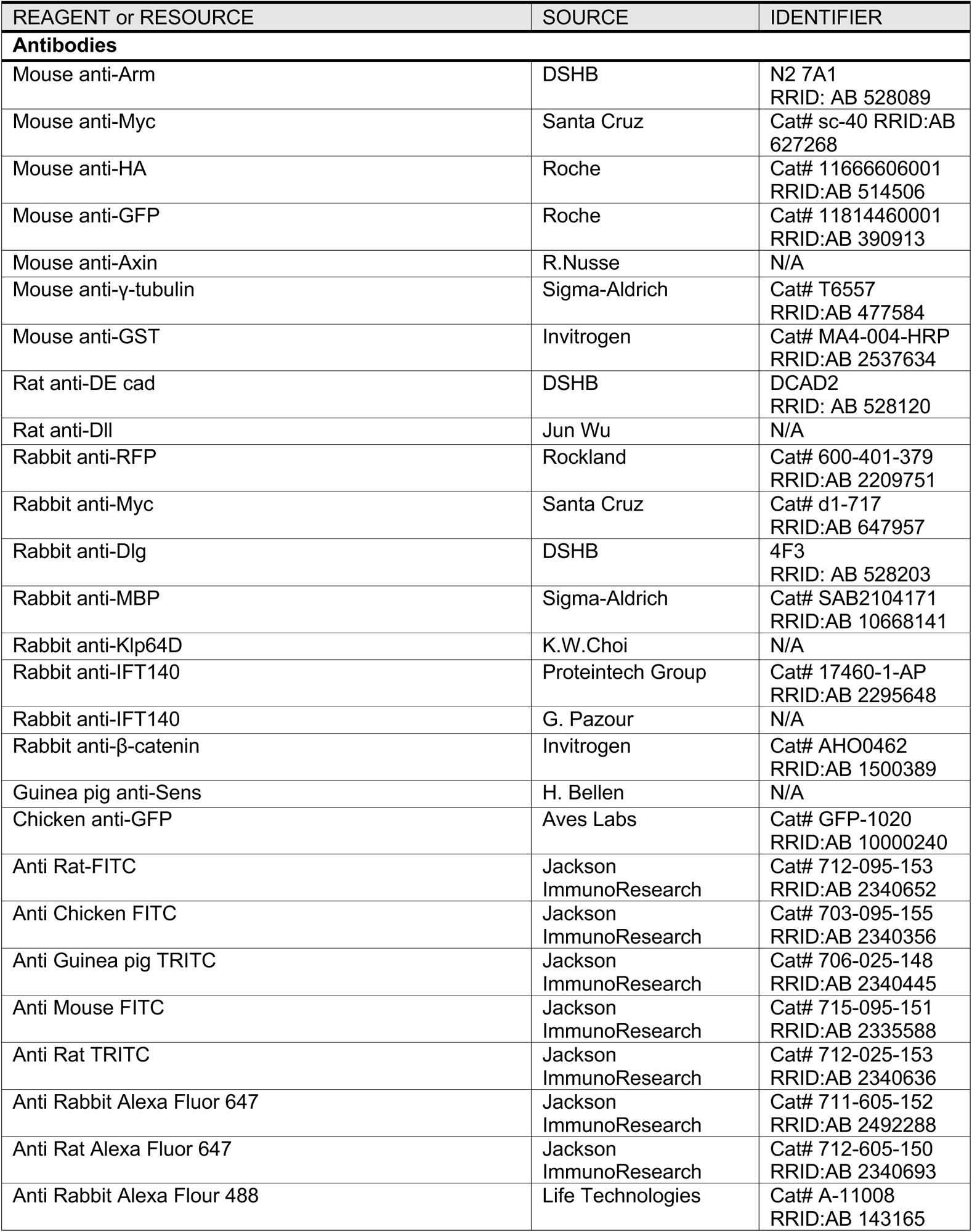
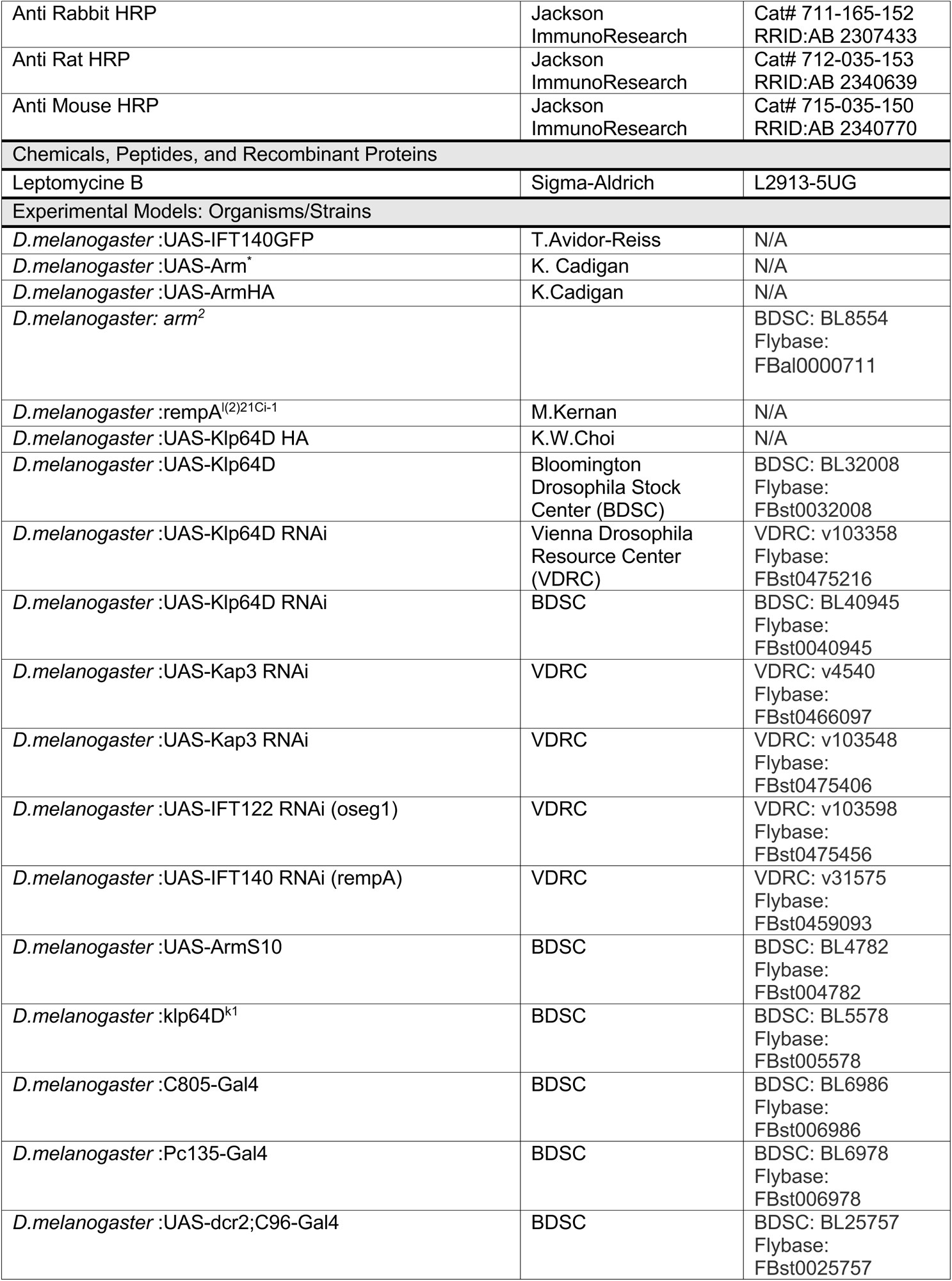
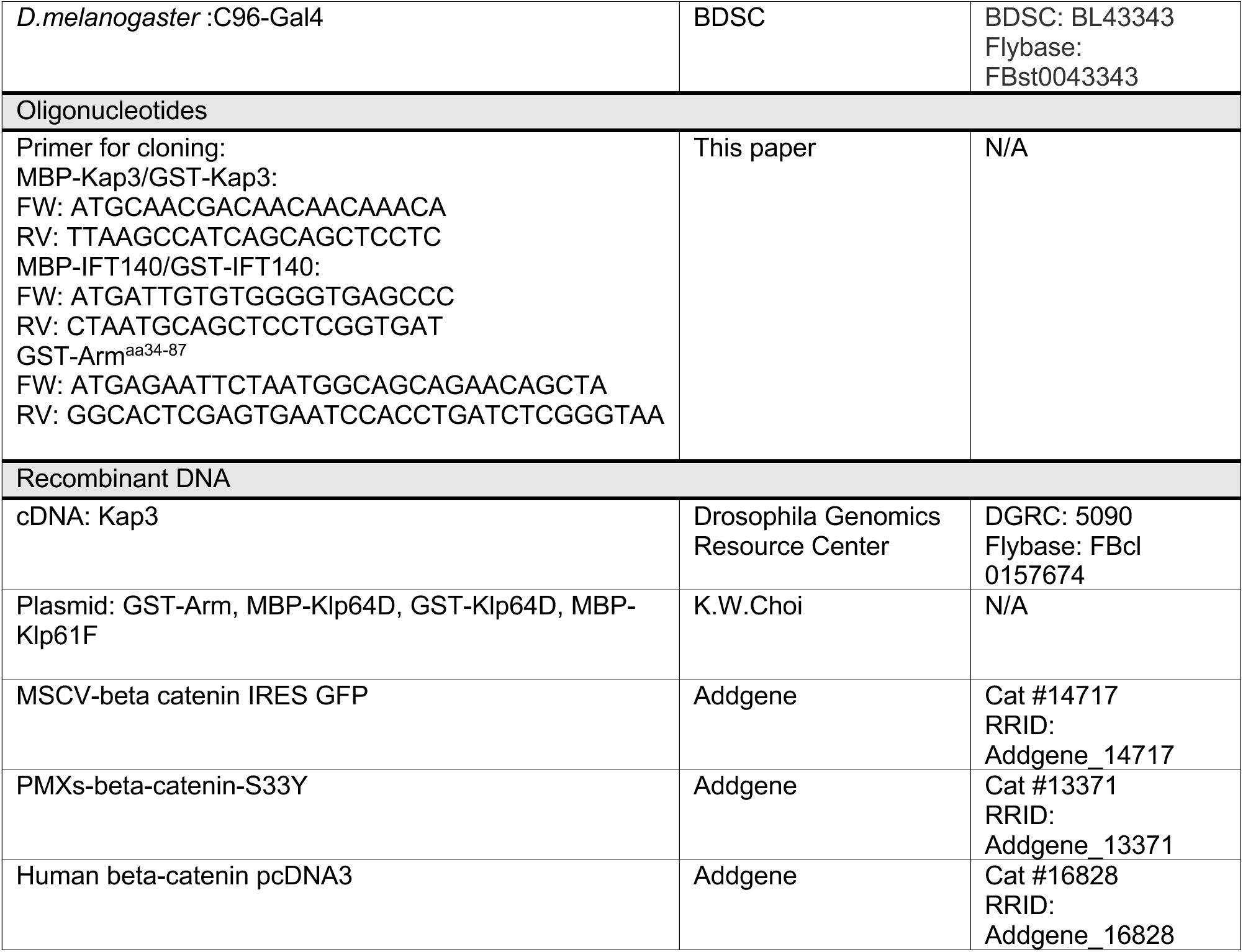

## ACKNOWLEDGEMENTS

We are grateful to Carlo Iomini, Davide Esposito, and Stuart A. Aaronson for reagents and advice, and Bo Chen for allowing us to use their tissue culture facility. We thank all Mlodzik lab members for helpful suggestions, Prashanth Rangan and Sam Sidi for helpful suggestions on the manuscript. We also thank the Bloomington Drosophila Stock Center for fly strains, and the Developmental Studies Hybridoma Bank at University of Iowa for antibodies. We would also like to thank the ISMMS Microscopy CoRE, where confocal microscopy was performed, which was in part supported by the Tisch Cancer Institute P30 CA196521 grant from the NCI. This work was supported by National Institutes of Health grant R35 GM127103 to MM and New York State Stem Cell (NYSTEM) science training award (postdoctoral fellowship to LTV).

## AUTHOR CONTRIBUTIONS

LTV and MM designed the study, LTV performed all experiments and developed experimental tools. MM and LTV analyzed the data and wrote the manuscript. MM provided funding for the study.

