## Supplemental Data for "Wg/Wnt-signaling induced nuclear translocation of β-catenin is attenuated by a β-catenin peptide through its interaction with IFT-A in development and cancer cells"

**Supplemental Figures**

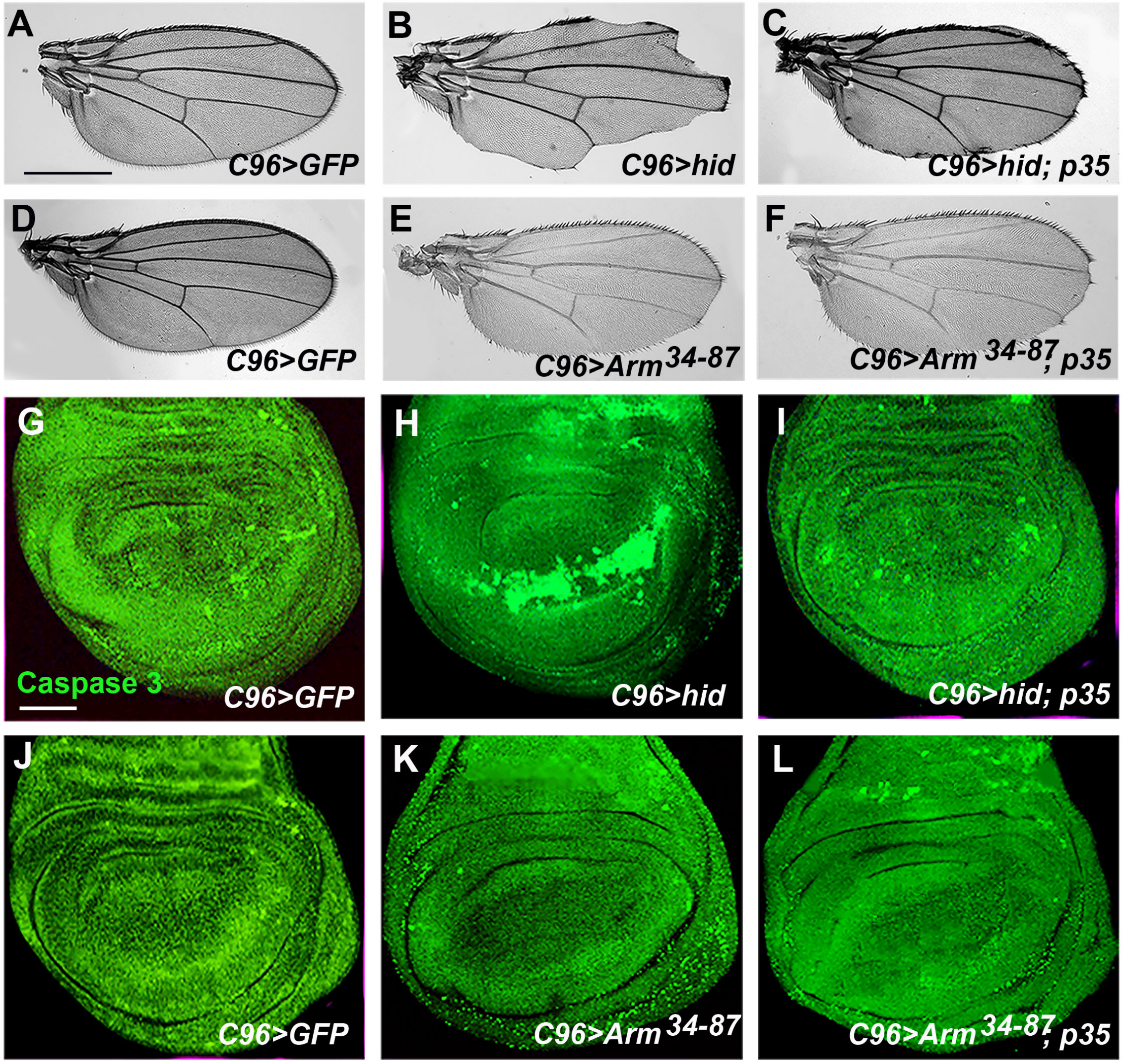


**Figure S1: Expression of the Arm$^{34\text{-}87}$ peptide does not induce cell death.**

(**A**) *C96>GFP* as a control. (**B**) Expression of Hid causes loss of wing margin. (**C**) Co-expression of Hid with p35 largely rescued the wing margin loss phenotype.

(**D**) Control wing (*C96>GFP*). (**E**) Wing notching by Arm$^{34\text{-}87}$ expression. (**F**) The Arm$^{34\text{-}87}$ effect is not suppressed by p35 co-expression.

(**G-I**) Hid expression by *C96-Gal4* induces caspase activation marked by cleaved Caspase 3 staining (**H**; compare to *C96>GFP* as a control, **G**). Caspase 3 detection (cell death) induced by Hid is suppressed by p35 co-expression (**I**).

(**J-L**) Arm$^{34\text{-}87}$ expression does not induce cell death (**K**, cf to control in **J**). p35 co-expression has no effect on Caspase 3 staining (**L**). Scale bar represents 100μm in A-F and 50μm in G-L.

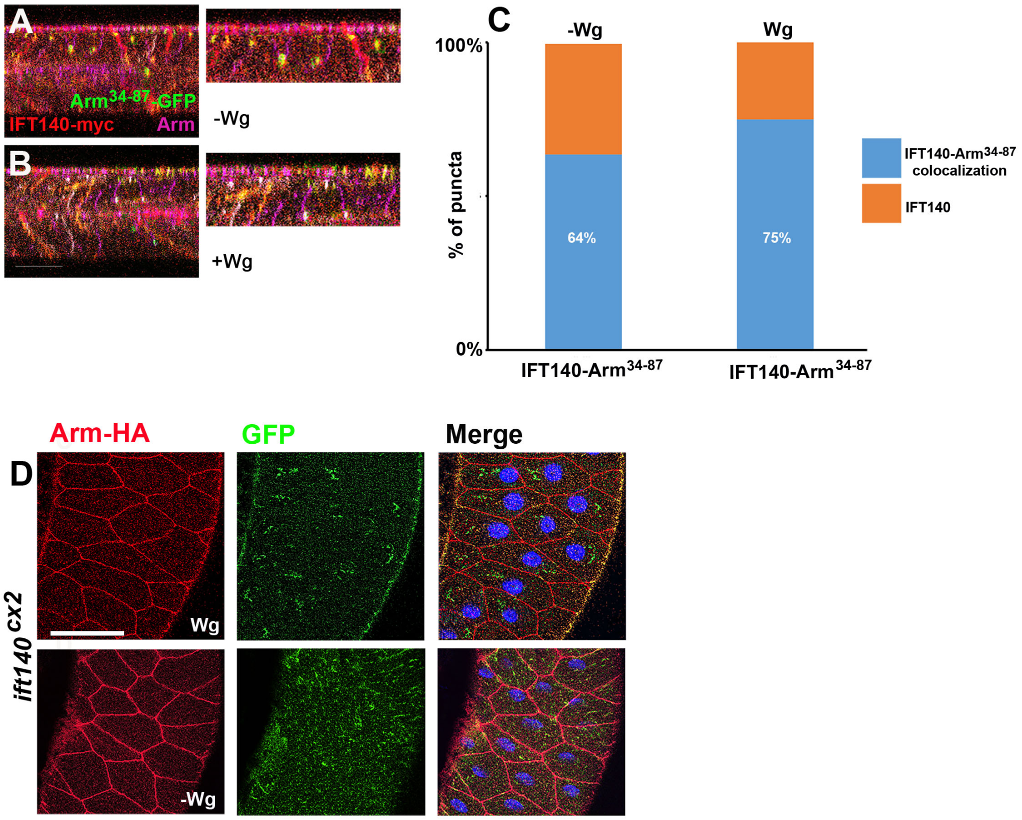


**Figure S2: Co-localization between IFT140, Arm and Arm$^{34\text{-}87}$ in *Drosophila* salivary gland cells.**

Salivary glands were stained for Arm$^{34\text{-}87}$-GFP (green), IFT140-myc (red), and Arm (magenta). (**A-B**) Z sections of salivary gland tissue. Note punctate staining displaying overlap at the sub-apical region of the salivary gland cells. (**A**) Without expression of Wg, endogenous Arm/β-catenin mainly localizes to the junctional regions of the membrane. Note punctate staining of Arm$^{34\text{-}87}$-GFP overlapping with IFT140-myc. (**B**) Upon Wg expression, Arm/β-catenin is stabilized and triple positive puncta, staining for Arm$^{34\text{-}87}$-GFP, IFT140-myc, and endogenous Arm are detected in the cytoplasm. (**C**) Quantification of the number of punctate containing IFT140 alone (orange) or Arm$^{34\text{-}87}$ with IFT140 (blue) (n>150 punctae per genotype from five different salivary glands). Note that Arm$^{34\text{-}87}$ and IFT140 co=localization does not depend on Wg-signaling.

(**D**) Salivary glands mutant for *ift140$^{cx2}$* were stained with Arm-HA (red), Arm$^{34\text{-}87}$-GFP (green), and Hoechst (blue). Note that the co-localization of full-length Arm and Arm$^{34\text{-}87}$-GFP, as induced by Wg expression (seen above in panel B, and Figure 2A-B main text), is lost in the *ift140$^{cx2}$* background. This suggests that wild-type IFT-A complexes can form multimers of unknown stoichiometry.  Note also that full-length Arm is not detected in the cytoplasm upon Wg-induction in the absence of IFT-A. Scale bar represents 50μm.

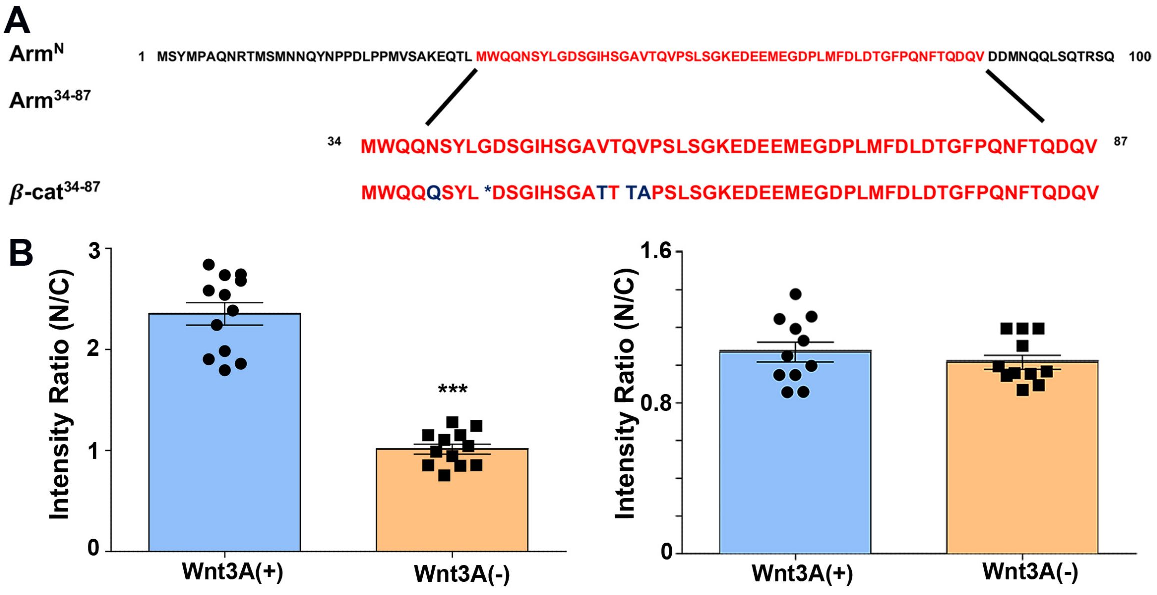


**Figure S3: Quantification of nuclear β-catenin and β-catenin$^{34-87}$-GFP in MEFs**

(A) Arm N terminal (Arm$^{N}$) and Arm$^{34-87}$ amino acids sequence. Note that the Arm$^{34-87}$/β-cat$^{34-87}$ region is highly conserved from *Drosophila* to higher animal. (B) Quantification of nuclear β-catenin-GFP and β-catenin$^{34-87}$-GFP immunofluorescence signal in MEFs as ration of nuclear (N) vs cytoplasmic (C) signal. Y axis denominates the ratio of green intensity values of selected regions (area of 4.705µm x 4.843µm) within nuclei and cytoplasm of individual cells (membrane-associated β-catenin was purposely excluded). Mean $\pm$ s.d. of values, obtained in randomly selected cells (*n*), are shown from five independent experiments; Student's t-test: *** $p$ <0.001; *ns*: difference statistically not significant.

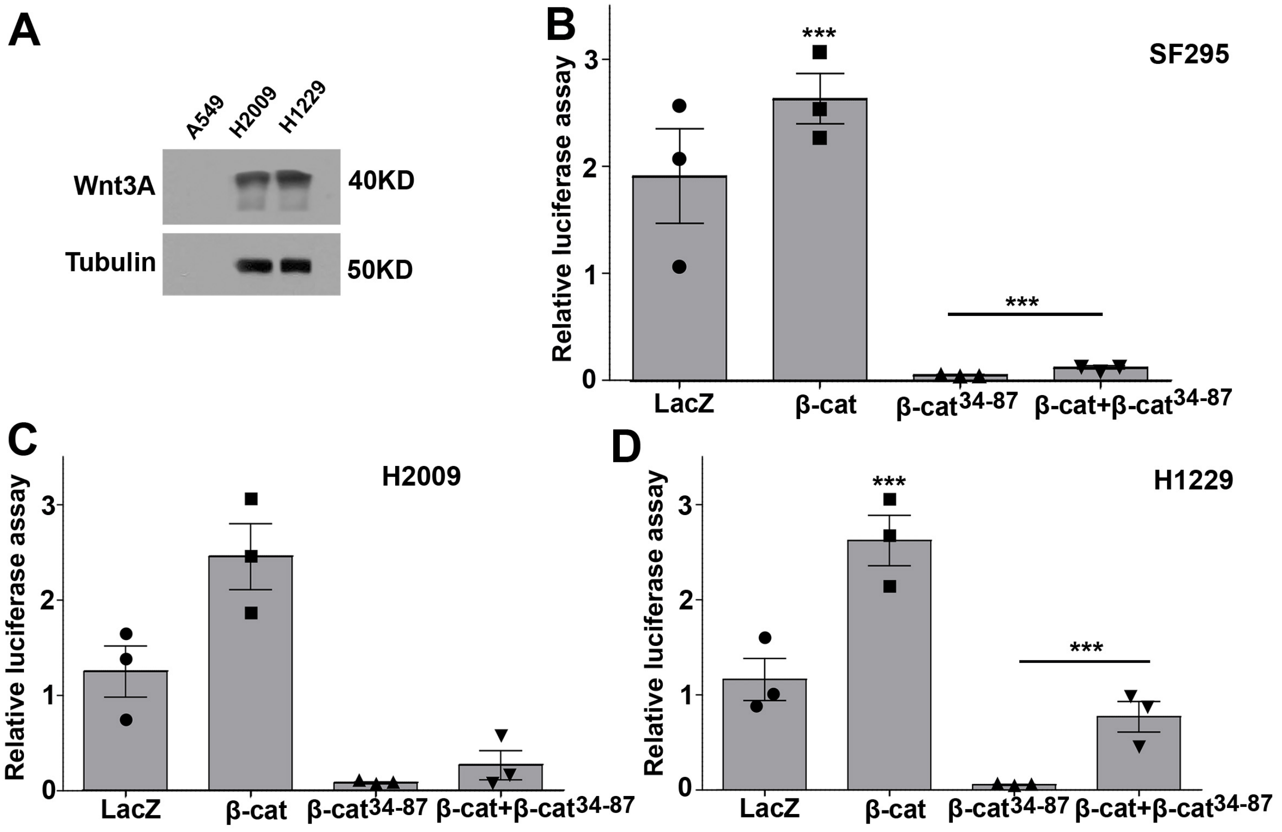


**Figure S4. The *β*-catenin$^{34\text{-}87}$ peptide strongly inhibits Wnt signaling in human cancer cells.**

(**A**) Detection of endogenous Wnt3A expression in lung cancer cell lines (A549, H2009 and H1229). Note high levels of Wnt3a in H2009 and H1229.

(**B-D**) Assay for Wnt-signaling activity in SF295 brain tumor cell line (B) and the two lung cancer lines H2009, and H1229 (C, D) transfected with either control LacZ, β-catenin, β-catenin$^{34\text{-}87}$ peptide, or β-catenin plus β-catenin$^{34\text{-}87}$ peptide, respectively. The relative luciferase activity indicates the ratio of TOP-Flash Wnt-reporter firefly luciferase and renilla luciferase activities. Overexpression of β-catenin increases the relative luciferase activity, whereas transfection with the β-catenin$^{34\text{-}87}$ peptide, or β-catenin plus β-catenin$^{34\text{-}87}$ peptide causes a strong reduction of Wnt-signaling activity in all tumor cells (***$p$ <0.001, three independent assays, student's t-test).
